# A Conserved Mechanism for Dimerization and Activation of Superfamily 1A UvrD-family Helicases

**DOI:** 10.64898/2026.05.20.726581

**Authors:** Binh Nguyen, Kacey N. Mersch, Ankita Chadda, Eric A. Galburt, Timothy M. Lohman

## Abstract

DNA helicases are ATP-dependent motor proteins that catalyze duplex DNA unwinding and are involved in DNA repair, recombination and replication restart. Prominent members of the non-hexameric SF1A UvrD-family helicases are *E. coli* UvrD, Rep, *B. stearothermophilus* PcrA and *M. tuberculosis* UvrD1. SF1A monomers are processive 3’ to 5’ single stranded DNA translocases, but need to be activated to become DNA helicases. One mechanism of activation is dimerization. Whereas Rep, UvrD and PcrA form non-covalent dimers, the *Mtb* UvrD1 helicase forms a redox-dependent covalent dimer. Dimerization of *Mtb* UvrD1 occurs between the same regulatory domain (2B) within each subunit stabilized by a disulfide bond formed between the same cysteine (Cys451) within each subunit. Dimerization relieves an inhibitory interaction between the 2B domain and duplex DNA within the monomer-DNA complex. We show here that Rep, UvrD and PcrA dimerize using the same 2B-2B interface. By placing a Cys residue within the 2B domains of Rep, UvrD and PcrA in the structurally equivalent position occupied by Cys451 of *Mtb* UvrD1, all three enzymes form redox-dependent covalent dimers that are constitutively active helicases with increased processivity compared to the non-covalent dimers. Hence, the 2B domain is a general dimerization domain for UvrD-family SF1A helicases.

## INTRODUCTION

Helicases are ATP-dependent enzymes that function in all aspects of DNA and RNA metabolism in all organisms (1-5). Helicases have been grouped into six superfamilies (SF) (6) and consist of two types depending on the oligomeric state of the functional helicase. SF3-6 are hexameric enzymes that have a toroidal structure that encircles one or both strands of DNA (7,8). SF1 and SF2 helicases are non-hexameric with functional forms that are generally monomeric or dimeric (9,10), although the oligomeric states of these enzymes have often not been characterized. Based on more stringent sequence homology, the SF1 DNA helicases can be further sub-divided into three groups (10), which are named after the founding members, UvrD-like, Pif1-like and Upf1-like. A further classification involves the directionality of translocation along single stranded (ss) nucleic acids, with SF1A enzymes (UvrD-like) translocating in the 3’ to 5’ direction, while SF1B enzymes (Pif1-like and Upf1-like) translocate in the 5’ to 3’ direction (7).

Some of the best studied members of the UvrD-family of SF1A DNA helicases are *E. coli* (*Ec*) UvrD, Rep, RecB, *B. stearothermophilus* (*Bst*) PcrA, *S. cerevisiae* Srs2, and *M. tuberculosis* (*Mtb*) UvrD1. Monomers of these enzymes share a common structure, shown in **Figure 1A**, with four sub-domains (11-15). The 1A and 2A sub-domains comprise the RecA motor domains, while the 1B and 2B sub-domains are accessory domains. The 2B sub-domain can undergo large rotations (as much as 160^°^ for UvrD) about a hinge connecting it to the 2A sub-domain (11-13,16-18). One important aspect of the UvrD-family of DNA helicases is that monomers can use ATP to translocate directionally (3’ to 5’) along ssDNA (19-28), but monomers show no processive helicase activity (27,29-31). Activation of SF1A helicase activity (9,32) occurs by several mechanisms, including monomer interaction with an accessory protein (MutL for UvrD (33,34), PriC for Rep (35) and RepD for PcrA (36)), removal of the 2B sub-domain (26,37), or formation of homo-dimers (18,27,29,31,38-41).

**Figure 1.**
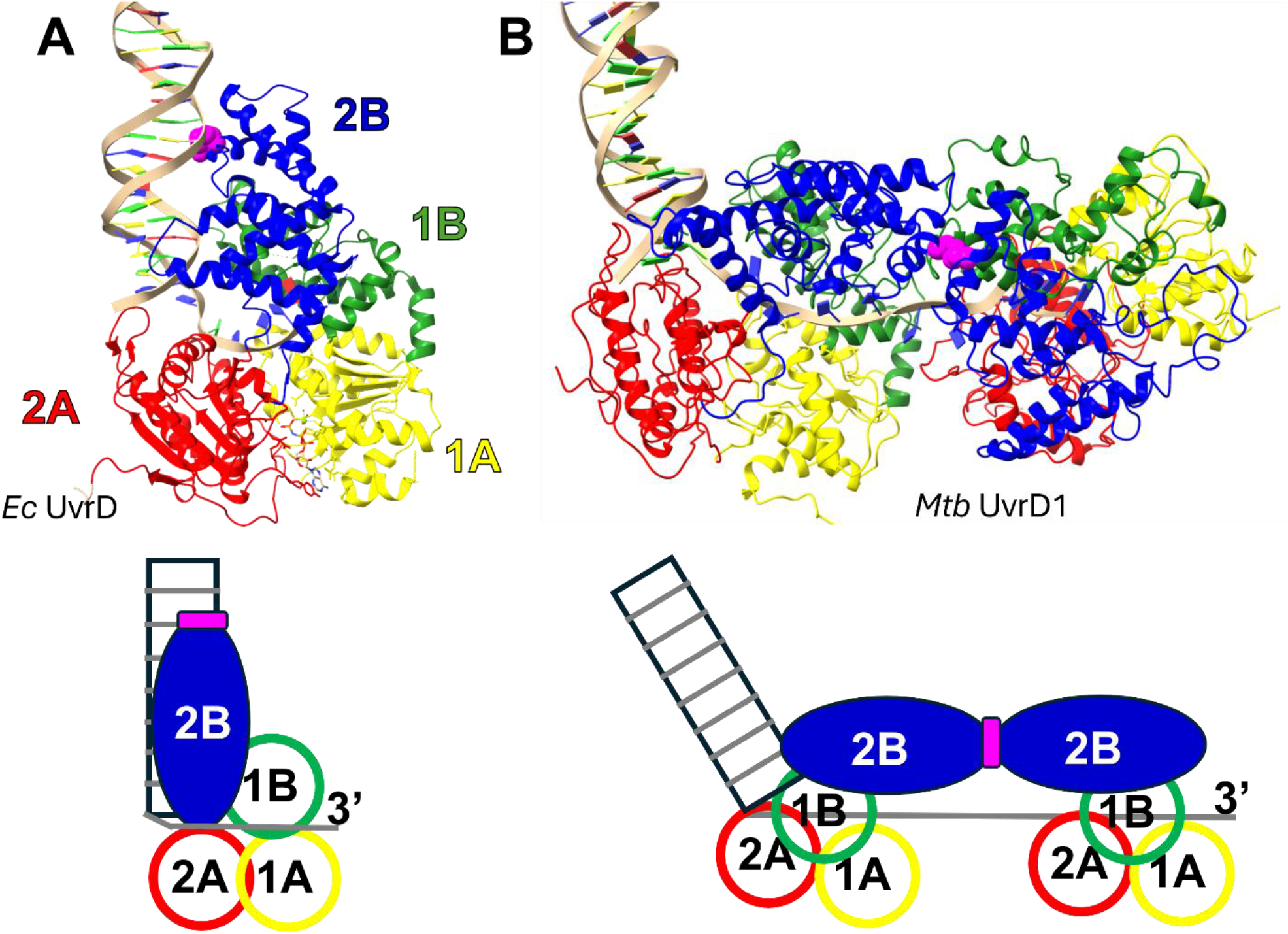
Structures of Monomeric UvrD and Dimeric UvrD1 bound to DNA junctions. (A)- Structure of *E. coli* UvrD∆40 monomer bound to a 18 base pair duplex with a 3’-(dN)_7_ single stranded DNA (PDB ID: 2IS4) (12). The four sub-domains are 1A (yellow), 2A (red), 1B (green) and 2B (blue). The R421 amino acid is shown in magenta. A cartoon showing that the 2B sub-domain interacts with duplex DNA in the monomer structure is shown below. (B)- Structure of dimeric *M. tuberculosis* UvrD1 bound to a duplex-(dT)_20_ junction (PDB ID: 9DES). The C451-C451 disulfide crosslinking the two 2B sub-domains (blue) is shown in magenta. A cartoon depicting the dimer-DNA complex is shown below.

Although evidence for helicase activation by dimerization has been reported for over 30 years, the mechanism and the identity of the dimerization interface has only recently been discovered in the context of *Mtb* UvrD1 (14,32). The *Mtb* UvrD1 dimer forms a redox-dependent dimer, stabilized by a disulfide bond formed between the same cysteine (Cys451) within the 2B sub-domain of each subunit (14,32) (**Figure 1B**). Previous crystal structures of *Ec* UvrD monomer (12) and *Bst* PcrA monomer (17) bound to a ss/duplex DNA junction show the 2B sub-domain in contact with the duplex DNA, while the 1A and 2A motor domains contact the ssDNA (**Figure 1A**). The same configuration involving the 2B sub-domain interaction with duplex DNA is observed for a *Mtb* UvrD1 monomer bound to a ss/duplex DNA junction (14). However, upon *Mtb* UvrD1 dimerization, this duplex DNA interaction with the 2B sub-domain is eliminated. In fact, the same region of the 2B sub-domain that contacts the duplex DNA in the monomer forms the 2B-2B dimerization interface. Hence dimerization and 2B sub-domain-duplex DNA binding are mutually exclusive(14,32), as depicted in Figure 1B, and the 2B sub-domain interaction with the DNA duplex is likely the basis of the auto-inhibition of monomeric helicase activity (9,32).

In contrast to *Mtb* UvrD1, *Ec* UvrD, Rep and *Bst* PcrA form non-covalent dimers. However, a mutant of *Ec* UvrD (UvrD(R421C) in which a Cys is placed in the 2B sub-domain in the structurally equivalent position as Cys451 of *Mtb* UvrD1 can form redox-dependent covalent dimers with constitutive helicase activity (14). This indicates that *Ec* UvrD dimers use the same 2B-2B interface as UvrD1 and suggests that the 2B sub-domain may be a general dimerization interface for UvrD-family SF1A helicases. Here we test this hypothesis further by making the same Cys mutation within the 2B sub-domains of *Ec* Rep and *Bst* PcrA. We find that this mutation enables both Rep and PcrA to form redox-dependent covalent dimers that are constitutively active helicases and conclude that the 2B sub-domain drives dimerization domain for this class of DNA helicases.

## MATERIAL AND METHODS

### Buffers and reagents

Buffer T is 10 mM Trizma base+HCl (pH 8.3), 20% (v/v) glycerol. Buffer A is Trizma base+HCl (pH 7.50 at 25 ºC), 10% (v/v) glycerol, 50 mM NaCl. Lysis buffer is 10 mM Trizma base+HCl (pH 8.3), 20% (v/v) glycerol, 10% (w/v) sucrose, 0.20 M NaCl, 0.20 mg/ml lysozyme, 1.0 mM Na_2_EDTA, 0.1 mM phenylmethylsulfonyl fluoride (PMSF). Storage buffer is 20 mM Trizma base+HCl (pH 8.3), 50% (v/v) glycerol, 0.20 M NaCl. ATP (Sigma-Aldrich) stock solutions were prepared in 50 mM NaOH (pH 7.5), and 200-μl aliquots were stored at -20 °C. ATP concentrations were determined spectrophotometrically using an extinction coefficient Ɛ_259_ = 15.4 × 10^3^ M^-1^ cm^-1^. MgCl_2_ concentrations were determined from refractive index measurements at 20 ºC with an ABBE Mark II refractometer using the reference values from the CRC handbook. DNA oligonucleotide sequences and their extinction coefficients are given in **Table S1**.

### Protein expression plasmidsThe plasmid expressing B

*stearothermophilus* PcrA mutant (S425C) was obtained from GENWIZ (Azenta Life Sciences). For PcrA(S425C), a missense mutation was introduced into pET-22b-pcrA, an ampicillin resistant plasmid containing the wild type PcrA gene (28), yielding a cysteine at position 425 of PcrA, (S425C) to create *pET-22b-pcrA(S425C)*. For Rep(A416C), a single mutation (A416C) was introduced into a plasmid expressing a cysteine-less Rep containing the following mutations (C18L, C43S, C167V, C178A, C612A)) (42) yielding the Kanamycin resistance plasmid (*pET28a-Rep(ΔCys, A416C)*). This plasmid has a 6x-His tag and a thrombin site at the N-terminus of Rep and confers kanamycin resistance. UvrD(R421C), was expressed from plasmid (*pA10-UvrD(ΔCys, R421C*). In UvrD(R421C) all 6 native cysteines were replaced with serine and a 6x-His tag and a thrombin cleavage site (MGSSHHHHHHSSGLVPRGSH) was placed at the N-terminus as described (16). A single mutation R421C was introduced into this plasmid to create (*pA10-UvrD(ΔCys, R421C)*).

### Protein purification

*E. coli* BL21(DE3) cells (Intact Genomics, St. Louis, MO, USA), transformed with the plasmid *pET-22b-pcrA(S425C)*, were grown in 5 liters of LB containing 50 µg/ml ampicillin (10-ml starter culture per liter) and induced with 0.5 mM IPTG at OD_600_ = 1.0 and grown for another 3 hours at 37 ºC. Cells were harvested by centrifugation at 4000xg for 20 minutes at 4 ºC. All purification steps were performed at 4 °C in buffer without reducing agents to preserve the dimer populations. Cell paste was resuspended in lysis buffer while stirring for 30 minutes. Lysate was sonicated on ice (Branson SFX250, Marshall Scientific, Hampton, NH, USA) until the solution was no longer viscous. The lysate was centrifuged at 30000xg for 90 minutes. The supernatant was extracted and ammonium sulfate (AS) powder was added to reach 41% saturation while stirring for 45 minutes. The protein was recovered in the AS pellet after centrifugation at 20000xg for 15 min. The AS pellet was resuspended in buffer T + 0.20 M NaCl and loaded onto a HiPrep Q HP 16/10 anion exchange column (Cytiva, Wilmington, DE, USA) (20-mL, preequilibrated with buffer T + 0.15 M NaCl) while mixing with buffer T without NaCl on the NGC chromatography system (Bio-Rad, Hercules, California, USA) to avoid precipitation (28). The column was washed with buffer T+ 0.15 M NaCl and eluted with a linear gradient from 0.15 to 0.50 M NaCl over 10 column volumes (cv). Fractions containing protein were run on two separate 12% acrylamide SDS gels with and without 2-mercaptoethanol to assess the fraction of monomers and dimers. Fractions containing monomers and dimers were loaded onto a 5-ml HiTrap Heparin HP Column (Cytiva) (28) at a conductivity equivalent to buffer + 0.15 M NaCl and eluted with a gradient from 0.15 M to 1.0 M NaCl over 40 cv. The monomer population elutes at a lower salt concentration than the dimers and fractions containing mostly dimers can be obtained. These fractions were then loaded onto a 9-ml ssDNA cellulose column equilibrated with buffer T + 0.10 M NaCl and protein was eluted with a NaCl gradient from 0.10 M to 2.0 M. To further purify the dimer population, the eluted protein was pooled, concentrated and loaded onto a 24-ml size-exclusion column Superdex 200 Increase 10/300 GL (Cytiva) with an 0.3-ml sample volume and a flowrate of 0.4 ml/min in buffer T + 0.5 M NaCl. The dimer fractions were pooled and dialyzed vs. storage buffer, flash-frozen with liquid nitrogen and stored at -80 ºC. The protein concentration was determined spectrophotometrically in 20 mM Tris (pH 7.5 at 25 °C) using ϵ_280_ = 7.58×10^4^ M^-1^ (subunit) cm^-1^, calculated from its amino acid sequence using SEDNTERP (43).

The Rep(A416C) mutant was overexpressed in *E. coli* BL21(DE3) cells while the UvrD(R421C) mutant was overexpressed in *E. coli* BL21(DE3)*ΔUvrD* cells (16). Cells were grown in Terrific Broth (Research Products International) with 50 µg/ml kanamycin at 37 ºC until OD_600_ = 0.6, then cooled to 16 ºC. Protein expression was induced by adding ITPG (1.0 mM) followed by overnight growth at 16 ºC. The induced cells were harvested at 4000xg for 20 min. Cell paste was added to lysis buffer as described above, and the lysate was adjusted to 0.48M NaCl before sonication for 3 minutes and centrifugation at 30000xg. Solid AS was added to the supernatant to reach 40% saturation. The AS pellet was resuspended in buffer T + 0.20 M NaCl before loading onto a 10-ml Ni-NTA column (Marvelgent Biosciences Inc, Waltham, MA USA), washed and eluted with buffer T + 0.20M NaCl + 0.50 M imidazole. The fractions from the Ni-NTA column containing protein were pooled and dialyzed (MWCO = 8k) against buffer T + 0.20 M NaCl to remove imidazole and the his-tag was cleaved with Thrombin (2 units of enzyme per mg protein). The proteins were subjected to Q-HP and Heparin HP columns as described above for PcrA. Fractions having mostly dimers were pooled and loaded on, dsDNA/ssDNA columns and eluted with buffer at 2.0 M NaCl as described (16,44). The eluted proteins were further subjected to a size exclusion chromatography using the Superdex 200 Increase 10/300 GL column as described above for PcrA. Fractions containing dimers were dialyzed against storage buffer, flash-frozen with liquid nitrogen and stored at -80 ºC. Protein concentrations were determined using the following extinction coefficients at 280 nm: 1.06×10^5^ M^-1^ (subunit) cm^-1^ for UvrD(R421C) (45) and 8.47×10^4^ M^-1^ (subunit) cm^-1^ for Rep(A416C) in 10 mM Tris (pH 7.5), 20% (v/v) glycerol, 0.10 M NaCl (46).

### Structural alignments

superposition of the 2B sub-domain structures of *Mtb* UvrD1, *Ec* UvrD, *Ec* Rep, and *Bst* PcrA were conducted using the pairwise alignment tool from the RCSB protein Data Bank (47). The 2B sub-domain of *Mtb* UvrD1 (PDB ID: 9DES, chain B, residues 404-589) was used as a reference structure to align *Ec* UvrD (PDB ID: 2IS4, chain A, residues 378-550), *Ec* Rep (PDB ID: 1UAA, chain A, residues 375-541), and *Bst* PcrA (PDB ID: 2PJR, chain A, residues 382-555). All structures were visualized using Chimera (48).

### Analytical ultracentrifugation

Sedimentation velocity (SV) experiments were conducted as described (27,35) with a ProteomeLab XL-A analytical ultracentrifuge (Beckman Coulter, Indianapolis, IN) at 25 ºC at 42000 rpm using a double-sector 12-mm Epon charcoal-filled centerpiece, and an 8-hole An-50Ti rotor. Protein concentrations were 100-200 nM and the cells scanned at 230 nm. UvrD(R421C) experiments were performed in buffer T + 20 mM NaCl. Rep(A416C) and PcrA(S425C) experiments were performed in buffer A both in the presence and absence of 1.0 mM dithiothreitol (DTT). Data were analyzed using SEDFIT to obtain the sedimentation coefficient distribution, c(s) (49). The sedimentation coefficients at 25 ºC in buffer, s_25,b_, were converted to s_20,w_ values (20 ºC in water) using the buffer densities and viscosities from SEDNTERP (50).

### Fluorescence stopped-flow kinetics

DNA unwinding experiments were conducted with a SX-20 stopped-flow instrument (Applied Photophysics, Leatherhead, Surrey, UK) as described (34,35). Cy5 fluorescence was excited with a 625-nm LED and fluorescence emission was passed through a 665-nm long pass filter. DNA unwinding substrates (Integrated DNA Technologies, Inc., Coralville, IA, USA) have variable duplex DNA lengths (18, 21, 23, 25, 40, 50 base pairs), each with a 3’-(dT)_20_ tail, the double strand DNA was labeled at the blunt ends with Cy5 and a Black Hole Quencher (BHQ2). DNA sequences are given in **Table S1**. DNA and protein were pre-incubated in one syringe and reactions were initiated by rapid mixing with 2.0 mM ATP + 2 mM MgCl_2_, plus 2 µM Trap DNA (3’-dT_40_-10bp hairpin (HP65), **Table S1**) in the second syringe. Typical DNA substrate concentrations were 10 nM (after mixing) with varying protein concentrations as specified. The protein trap prevents any free protein from rebinding and reinitiating DNA unwinding, thus ensuring single round conditions. Fluorescence amplitudes were converted to fractions of DNA molecules unwound using the Cy5-labeled DNA only as a reference for 100% unwound DNA.

DNA unwinding time courses for different duplex lengths, L, plotted as the fraction of ssDNA molecules, *f*_*ss*_*(t)*, were fit to the n-step kinetic model in **Scheme (1)** using **Eq. (1)** (34,51).

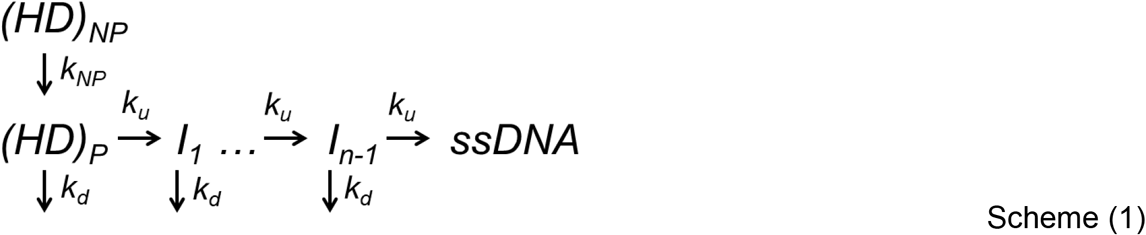

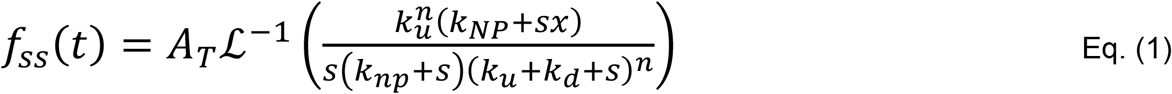

where A_T_ is the fraction of DNA molecules unwound; k_u_ is the unwinding rate constant per step and k_d_ is the dissociation rate constant; n is the number of steps; k_NP_ is the rate from non-productively bound (NP) complex to productively bound complex, and *x* is the fraction of productively bound complex ((HD)_P_ / ((HD)_P_ + (HD)_NP_)). ℒ^-1^ is the inverse Laplace transform operator, and s is the Laplace variable. The DNA unwinding step size, m = L/n (bp/step). The macroscopic unwinding rate (bp/s) is mk_obs_, where k_obs_ = (k_u_ + k_d_), and k_obs_ ≈ k_u_ when k_u_ >> k_d_. The DNA unwinding time courses were well described by Scheme 1. A_T_ is related to the processivity P as in Eq. (2).

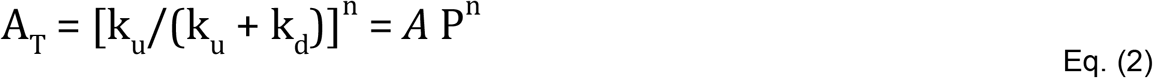

The average number of base pairs unwound per binding event, <N_bp_> was obtained from fitting A_T_ as a function of duplex length, L, using Eq. (3).

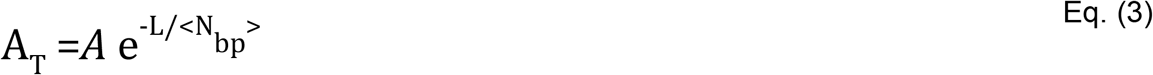

## RESULTS

*Formation of redox-dependent cross-linked dimers of Ec UvrD, Ec Rep, and Bst PcrA. Mtb* UvrD1 helicase requires dimerization in order to activate the helicase (27). Subsequent cryo-EM structures of the *Mtb* UvrD1 dimer identified the 2B sub-domains as the primary dimerization interface, stabilized by a disulfide bridge between Cys451 in each 2B sub-domain (14). *Ec* UvrD (30,31,38,52), *Ec* Rep (26,29,37) and *Bst* PcrA (28) require non-covalent dimerization for helicase activation in the absence of accessory factors or force. As such, we sought to determine whether the same 2B sub-domain interface is used to facilitate dimerization in those enzymes. **Figure 2A** shows the region of the 2B sub-domain of *Mtb* UvrD1 containing Cys451 aligned with the same regions of the 2B sub-domains of *Ec* UvrD, *Ec* Rep and *Bst* PcrA. In the sequence alignment, *Mtb* Cys451 aligns with *Ec* UvrD V425. However, Chadda et al. (14) showed that a V425C mutation in *Ec* UvrD did not form a disulfide linked UvrD dimer. Superposition of the structures of the 2B sub-domains of *Mtb* UvrD1 with *Ec* UvrD, indicated that R421 of *Ec* UvrD was in the structurally equivalent position to *Mtb* UvrD1 C451. In fact, the *Ec* UvrD (R421C) mutation readily formed a redox-dependent dimer that possessed constitutive helicase activity even under conditions of excess DNA which promotes UvrD monomer-DNA complexes for wild type *Ec* UvrD. Superposition of the structures of the 2B sub-domains of Rep and the PcrA with the *Mtb* UvrD1 2B sub-domain (**Figures 2C** and **D**), show that Rep A416 and PcrA S425 superimpose with *Ec* UvrD R421 and *Mtb* UvrD1 C451. In fact, UvrD R421, Rep A416 and PcrA S425 also align in the sequence (**Figure 2A**).

**Figure 2.**
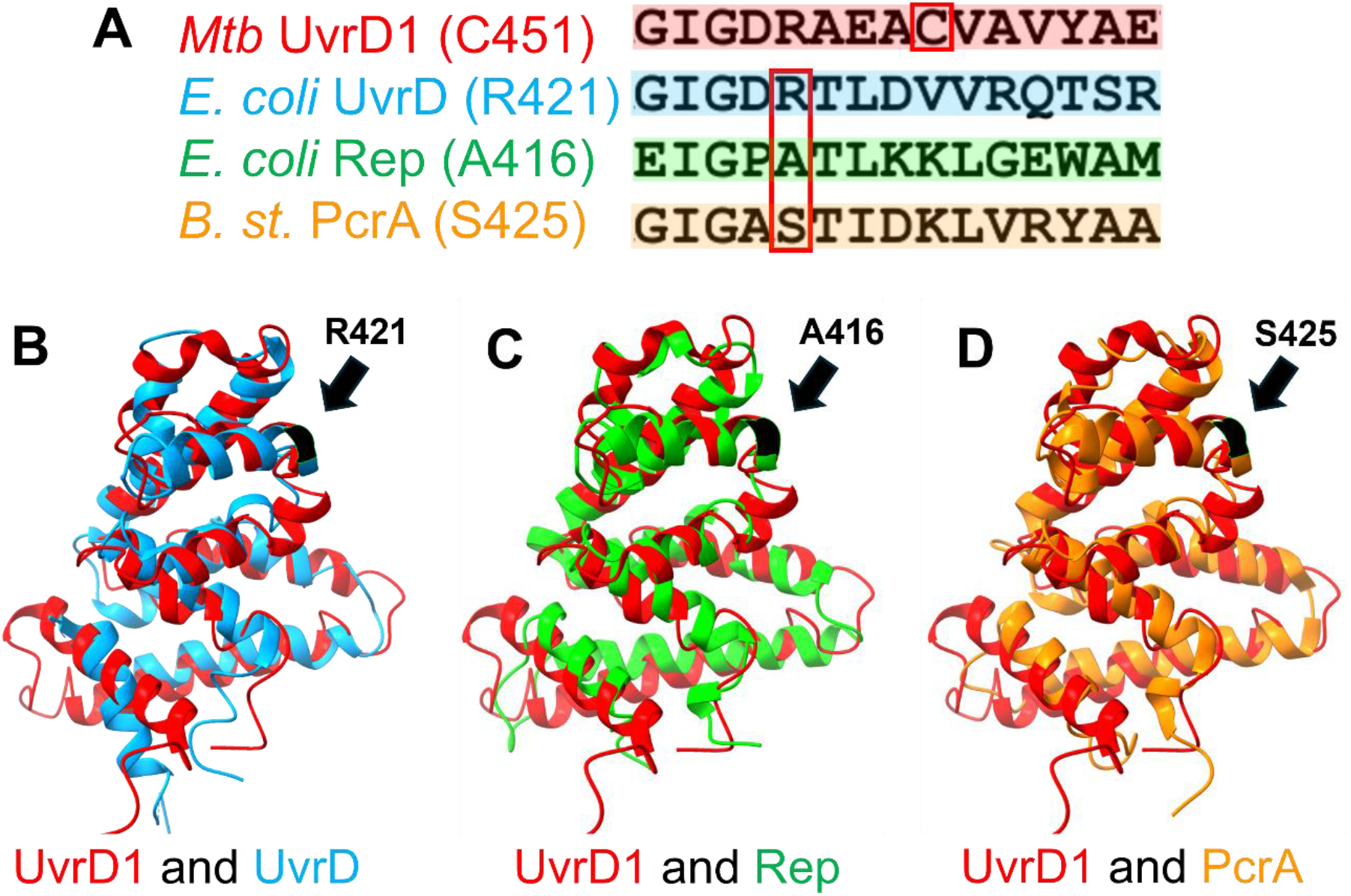
Superposition of the 2B sub-domains of UvrD1, UvrD, Rep and PcrA. (A)- Sequence alignment of the region of the 2B sub-domain of *Mtb* UvrD1 that contains Cys 451 with the same regions of *E. coli* UvrD, *E. coli* Rep and *B. stearothermophilus* PcrA. The amino acids that were changed to Cys are boxed in red. (B)- Superposition of the 2B sub-domain of *Mtb* UvrD1 (red) and *E. coli* UvrD (blue) showing that UvrD R421 is in the equivalent position of *Mtb* UvrD1 C451. (C)- Superposition of the 2B sub-domain of *Mtb* UvrD1 (red) and *E. coli* Rep (green) showing that Rep A416 is in the equivalent position of *Mtb* UvrD1 C451. (D)- Superposition of the 2B sub-domain of *Mtb* UvrD1 (red) and *Bst* PcrA (orange) showing that PcrA S425 is in the equivalent position of *Mtb* UvrD1 C451.

After purification in the absence of reducing agents (Methods), sedimentation velocity experiments revealed that Rep(A416C), PcrA(S425C), *Bst* PcrA(S425C) exist in both monomeric and dimeric species **(Figure 3A-C)**. In each case, two peaks are observed, corresponding to monomers (M) with s_20,w_= 4.6-4.8 S and disulfide cross-linked dimers (D) with s_20,w_=7.0-7.5 S. The extent of dimer formation was highest for PcrA(S425C) (∼80%), less for UvrD(R421C) (∼35%) and lowest for Rep(A416C) (∼27%). Subsequent size-exclusion chromatography (Materials and Methods) was used to isolate the dimer fraction (**Figure 3 (D-F**) and addition of 1 mM DTT completely eliminates dimerization **(Figure 3G-I)** confirming that dimerization is redox dependent as is the case for *Mtb* UvrD1 (27). The low yield of Rep(A416C) dimers is consistent with previous observations that wild type Rep dimerization is DNA-dependent (40,41,53).

**Figure 3.**
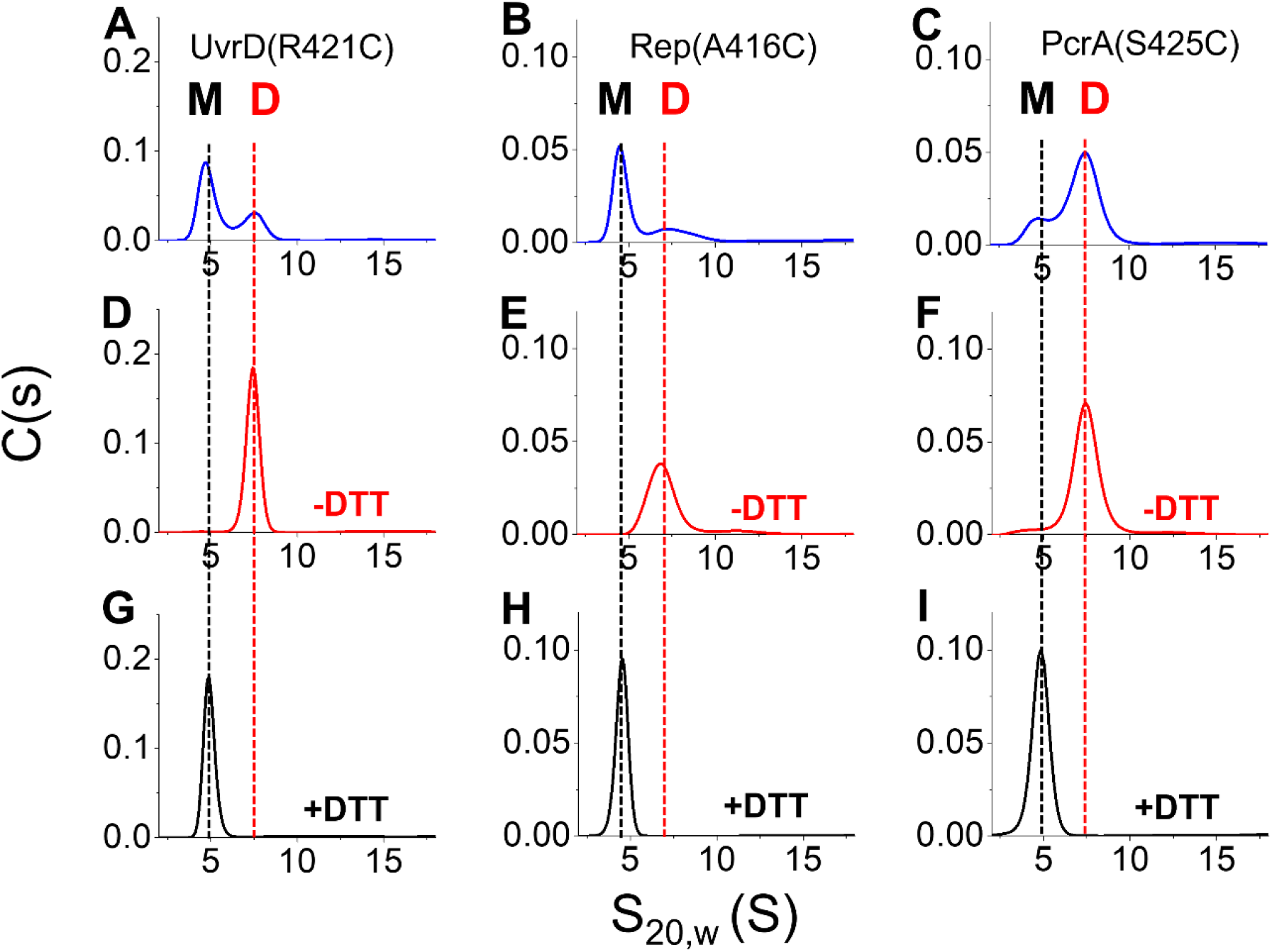
*E. coli* UvrD (R421C), *E. coli* Rep (A416C) and *Bst* PcrA (S425C) form disulfide crosslinked dimers under oxidizing conditions. (A-C)- Sedimentation velocity profiles (-DTT) of *E. coli* UvrD(R421C), *E. coli* Rep(A416C) and *Bst* PcrA(S425C) purified in the absence of reducing agent show a mixture of monomers (UvrD(R421C) M, s_20,w_=4.8 S and covalent dimers D (35%), s_20,w_=7.5 S; Rep(A416C) M, s_20,w_=4.6 S and D (27%), s_20,w_=7.0 S; PcrA(S425C) M, s_20,w_=4.8 S and D (80%), s_20,w_=7.4 S). (D-F)- Sedimentation velocity profiles (-DTT) of purified dimers after gel filtration to separate the dimers from the monomers. (G-I)- Sedimentation velocity profiles showing that each protein transitions to only stable monomers under reducing conditions (+ 1.0 mM DTT). Experiments were conducted at 25 ºC in buffer T+ 20 mM NaCl for UvrD(R421C) and in buffer A for Rep(A416C) and PcrA(S425C). The protein concentrations (in monomer units) were 0.10 µM (UvrD(R421C) and Rep(A416C)) and 0.20 µM for PcrA(S425C).

*The crosslinked dimers are constitutively active helicases*. We used a fluorescence stopped-flow assay to examine the DNA helicase activity of the covalent UvrD, Rep and PcrA dimers as described (34). The DNA molecules used are depicted schematically in **Figure 4A** and consist of an 18 base pair duplex with a 3’-(dT)_20_ flanking ssDNA. The blunt ends of the DNA duplex are 5’-labeled with a Cy5 fluorophore and 3’-labeled with a black hole quencher (BHQ2). The BHQ2 quenches the Cy5 fluorescence within the duplex and DNA unwinding is monitored by the increase in Cy5 fluorescence upon strand separation. Protein and DNA substrates are pre-mixed in one syringe of the stopped-flow and the reaction is initiated by mixing with buffer containing ATP (1.0 mM) and a trap for free protein to prevent re-initiation and thus ensure a single round of DNA unwinding.

**Figure 4.**
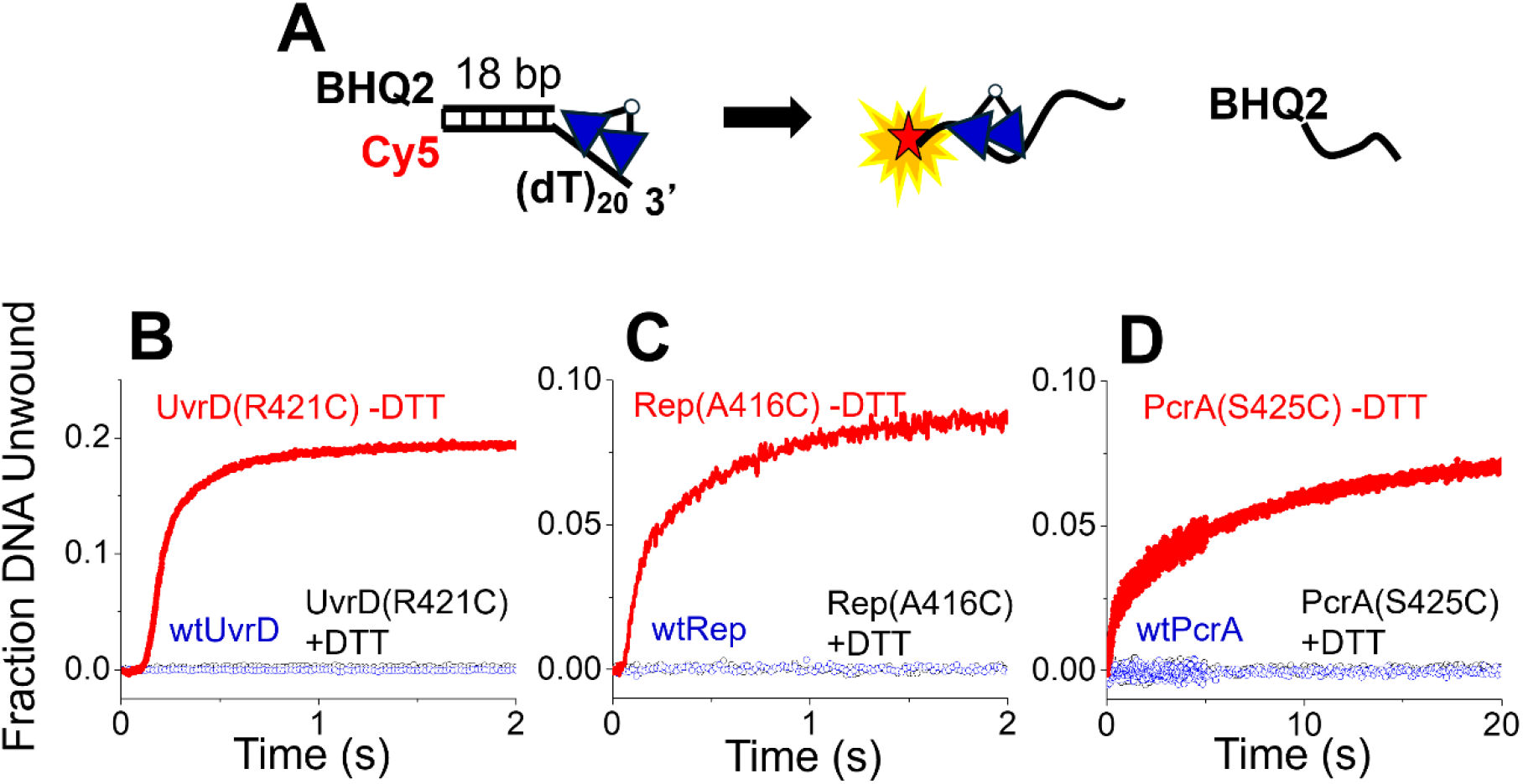
Covalent dimers of *E. coli* UvrD(R421C), *E. coli* Rep(A416C) and *Bst* PcrA(S425C) show constitutive helicase activity. (A)- DNA unwinding of an 18 base pair duplex with a 3’-(dT)_20_ flanking region was monitored by the increase in Cy5 fluorescence upon formation of ssDNA. (B-D)- Single round stopped-flow kinetic time courses catalyzed by 100% crosslinked dimers of (B)-UvrD(R421C); (C)- Rep(A416C) and (D)- PcrA(S425C) under conditions of 1 dimer per 2 DNA molecules (red traces -DTT-100% dimers), (black traces + 1.0 mM DTT-100% monomers). WtUvrD, wtRep, and wtPcrA show no unwinding at this 1 monomer : 1 DNA mixing, (blue traces). Experiments were conducted at 25 ºC in buffer T + 20 mM NaCl for UvrD(R421C) and in buffer A for Rep(A416C) and PcrA(S425C). The DNA and protein concentrations (in monomer units) after mixing were 10 nM at 1.0 mM ATP.

**Figure 4 (B-D)** shows the resulting kinetic time courses for unwinding of an 18 bp duplex by (**B)** UvrD(R421C) dimers, (**C**) Rep(A416C) dimers and (**D**) PcrA(S425C) dimers. All protein samples consisted of >98% covalent dimers and experiments were performed under conditions of excess DNA (1 dimer per 2 DNA molecules) at 25 ºC in buffer T+ 20 mM NaCl for UvrD(R421C) and in buffer A for Rep(A416C) and PcrA(S425C). The DNA and protein concentrations (in monomer units) after mixing were 10 nM. All three dimers showed significant DNA unwinding activity (red traces) under oxidative conditions (-DTT) that support dimerization, but no activity under reducing conditions (+DTT) that promote monomers. Under these conditions, the fraction of DNA molecules unwound was ∼20% for *Ec* UvrD(R421C), ∼8% for *Ec* Rep(A416C) and ∼7% for *Bst* PcrA(S425C). Although these extents of DNA unwinding may appear low, we note that the binding affinities of these dimers are not high enough to ensure that all of the dimers are bound to DNA at these enzyme and DNA concentrations (10 nM). At these same concentrations (10 nM), since an excess of wild type enzymes is needed to promote non-covalent dimerization and observe DNA unwinding.

*Crosslinked dimers unwind DNA at rates similar to those of wild type dimers but with higher processivity*. We next examined the rates and processivities of DNA unwinding by the crosslinked dimers by performing stopped-flow kinetics experiments with a series of fluorescent (Cy5/BHQ2) DNA substrates with different DNA duplex lengths (18, 21, 23, 25, 40 and 50 bp), all possessing a 3’-(dT)_20_ ssDNA flanking region.

Stopped-flow DNA unwinding was examined as described above with protein samples that were >98% dimer by pre-mixing protein and DNA in one syringe at a ratio of 1 dimer per 2 DNA. Reactions were initiated by mixing with buffer containing ATP. Final concentrations were 10 nM for DNA and 1.0 mM ATP. The resulting time courses for UvrD(R421C) (**Figure 5A**), and Rep(A416C) (**Figure 5B**), show distinct lag phases that increase with increasing DNA duplex length as expected for an all-or-none DNA unwinding assay in which the helicase proceeds through multiple steps with similar rate constants in order to fully unwind the DNA (51,54). The kinetic traces for PcrA(S425C) (**Figure 5C**) show much less of a lag phase, although a lag is still observed. Each series of kinetic traces was analyzed using a sequential “n-step” mechanism (black lines) (**Scheme 1, Eq. 1**, in Materials and Methods) to obtain a macroscopic rate of DNA unwinding of 71±2 bp/s for *Ec* UvrD(R421C), 138±20 bp/s for *Ec* Rep(A416C) and 73±10 bp/s for *Bst* PcrA(S425C) (see **Table 1**).

**Table 1.**
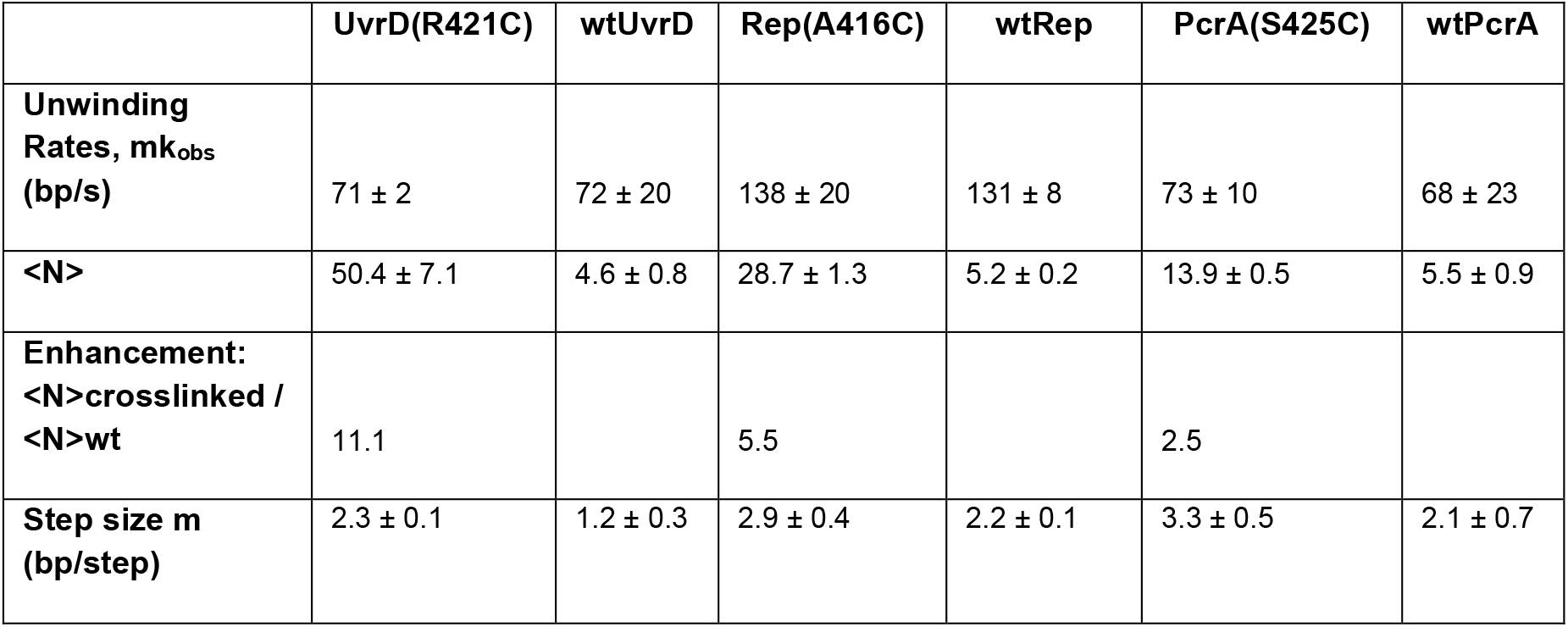
Kinetic parameters for DNA unwinding by crosslinked and wild type dimers using Scheme 1.

|  | UvrD(R421C) | wtUvrD | Rep(A416C) | wtRep | PcrA(S425C) | wtPcrA |
| --- | --- | --- | --- | --- | --- | --- |
| **Unwinding Rates, $m_{k_{obs}}$ (bp/s)** | 71 ± 2 | 72 ± 20 | 138 ± 20 | 131 ± 8 | 73 ± 10 | 68 ± 23 |
| **<N>** | 50.4 ± 7.1 | 4.6 ± 0.8 | 28.7 ± 1.3 | 5.2 ± 0.2 | 13.9 ± 0.5 | 5.5 ± 0.9 |
| **Enhancement: $\frac{\langle N \rangle_{crosslinked}}{\langle N \rangle_{wt}}$** | 11.1 |  | 5.5 |  | 2.5 |  |
| **Step size $m$ (bp/step)** | 2.3 ± 0.1 | 1.2 ± 0.3 | 2.9 ± 0.4 | 2.2 ± 0.1 | 3.3 ± 0.5 | 2.1 ± 0.7 |

**Figure 5.**
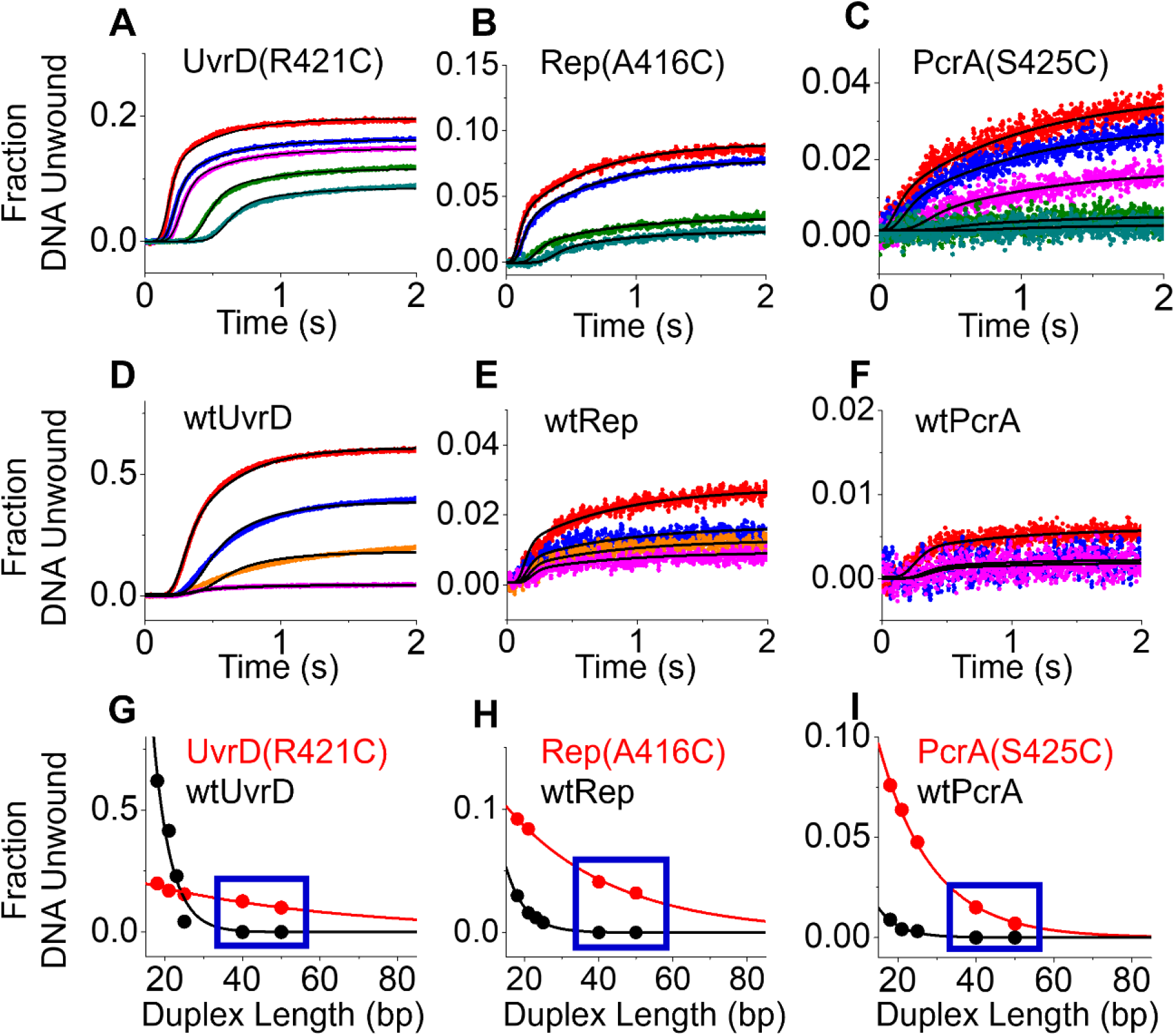
Covalent dimers of *E. coli* UvrD(R421C), *E. coli* Rep(A416C) and *Bst* PcrA(S425C) show higher DNA unwinding processivity than non-covalent wild type dimers. (A-C)- Single round stopped-flow DNA unwinding time courses using 100% dimer for a series of DNA molecules with a 3’-(dT)_20_ tail and varying duplex lengths (bp) (18 (red), 21 (blue), 23 (orange), 25 (magenta), 40 (olive), and 50 bp (dark cyan)) as indicated for (A)- UvrD(R421C); (B)- Rep(A416C) and (C)- PcrA(S425C) under excess DNA conditions (1 dimer/2 DNA). The DNA and protein concentrations (in monomer units) after mixing were 10 nM. (D-F)- Single round stopped-flow DNA unwinding time courses using wild type (un-crosslinked) enzyme for a series of DNA molecules with a 3’-(dT)_20_ tail and varying duplex lengths under excess protein conditions as indicated: (D)- wtUvrD (30 nM wtUvrD (monomer units) + 10 nM DNA); (E)- wtRep (100 nM wtRep (monomer units) + 10 nM DNA) and (F)- wtPcrA (80 nM wtPcrA (monomer units) + 10 nM DNA). Concentrations are after mixing. Black lines are fits using Eq. 1. (G-I)- Final DNA unwinding amplitudes for the kinetic traces in (A)-(F) for each duplex length. Boxed data indicate that the crosslinked dimers show significant DNA unwinding of 40 and 50 bp duplex DNA even in excess DNA, whereas the un-crosslinked wt enzymes show no DNA unwinding of 40 and 50 bp DNA even in 3-10-fold excess protein (monomer units).

For comparison, we also performed stopped-flow DNA unwinding experiments with each of the wild type enzymes at the same DNA concentration (10 nM) (**Figure 5 D-F**). However, since the wild type enzymes must dimerize in order to function as helicases, these experiments needed to be performed with enzyme in large excess over the DNA substrate. (3 wtUvrD per DNA; 10 wtRep per DNA; 8 wtPcrA per DNA). Within error, the DNA unwinding rates for the wild type non-covalent dimeric enzymes are the same as those for the cross-linked dimeric enzymes (see **Table 1**). However, as shown in **Figure 5 (G-I)** and **Table 1**, the processivities of DNA unwinding are significantly higher for the cross-linked dimers. We estimate that the processivity increases more than 11-fold for UvrD(R421C), five-fold for Rep(A416C), and two-fold for PcrA(S425C). The boxed data in **Figure 5 (G-I)** show that the crosslinked dimers can unwind DNA substrates that are 40 and 50 base pairs in length even in excess DNA (1 dimer / 2 DNA), whereas the wild type enzymes cannot unwind these duplex lengths even with protein in excess over DNA.

## DISCUSSION

Prior to the recent determination of the structure of the *Mtb* UvrD1 dimer (14), only monomeric structures had been reported for *Ec* UvrD (12), *Ec* Rep (13) and *Bst* PcrA (11,17). The structures of UvrD and PcrA monomers bound to ss/ds DNA junctions showed the 2B sub-domain in contact with the duplex DNA (12,17). This 2B-duplex DNA interaction was interpreted as being critical for promoting DNA unwinding (55,56). However, extensive evidence has shown that these enzymes need to be activated by dimerization in the absence of accessory proteins (9,32) or assisting force (57). Furthermore, deletion of the 2B sub-domain of Rep to form Rep∆2B activates the monomeric helicase indicating that the 2B sub-domain is an auto-inhibitory regulatory domain and not critical for DNA unwinding activity (26,37,58). However, the absence of dimer structures and the suggestion of unwinding in the context of the monomer structures (12,17) led to continued controversy. The cryo-EM structures of the *Mtb* UvrD1 dimer bound to a DNA junction under solution conditions that support helicase activity resolved this controversy (14,32). In the UvrD1 dimer structure, the dimer interface is formed between the 2B sub-domains of the two subunits and is stabilized by a disulfide crosslink formed between Cys451 in each of the 2B sub-domains (14,27). Importantly, dimerization prevents formation of the 2B sub-domain duplex DNA interactions that are observed in the monomeric UvrD and PcrA DNA structures (12,17). In fact, the monomeric *Mtb* UvrD1-DNA junction structures show the same 2B sub-domain-duplex DNA contacts (14). These observations argue that the 2B sub-domain-duplex DNA interaction is auto-inhibitory rather than catalytic and that dimerization activates the helicase by causing a large movement of the 2B sub-domain away from the duplex DNA thus relieving the inhibition (14,32) as shown in **Figure 1B**.

The question remained as to whether the 2B-2B interaction and the release of auto-inhibitory contacts is conserved across the non-covalent dimers formed by *Ec* UvrD, *Ec* Rep and *Bst* PcrA helicases. We showed previously that the *Ec* UvrD dimer was activated via the same 2B-2B dimerization interface and we show here that *Ec* Rep and *Bst* PcrA dimers are activated in the same manner. When a Cys residue is placed in the same position within the 2B sub-domain as found in *Mtb* UvrD1 (14), each enzyme can form a redox-dependent covalent dimer that is a constitutively active helicase. These covalent dimers have very similar DNA unwinding rates as the non-covalent dimers, but possess higher DNA unwinding processivities. The higher processivities likely reflect the fact that dimers have two subunits that each can bind DNA, hence if one subunit dissociates, the other can remain bound to the DNA.

The RecBCD and RecBC helicases are also stabilized by interactions between the 2B sub-domains of RecB, a UvrD-family SF1A helicase, and the 2B sub-domain of RecC (59), although those interactions use an interface that differs from that used by UvrD1, UvrD, Rep and PcrA dimers. The RecB monomer is a poor helicase(60), but is activated by interactions with RecC (25). We hypothesize that this activation likely involves a similar removal of the inhibitory RecB 2B sub-domain interactions with duplex DNA.

We emphasize that the 2B sub-domain of UvrD-family SF1A helicases/translocases has two functions depending on context. Within a dimer, it serves as the primary dimer interface and results in activation as this interaction sequesters the domain away from downstream duplex DNA (32). Within a monomer, the 2B sub-domain inhibits helicase activity by binding to duplex DNA and preventing DNA unwinding at a 3’ss/ds DNA junction. This inhibition at a 3’-ss/ds DNA junction may serve a biological function *in vivo* since monomeric *Ec* UvrD is a processive single stranded DNA translocase that functions to remove RecA filaments in its capacity as an anti-recombinase (61-63). *S. cerevisiae* Srs2 enzyme, another UvrD-family helicase/translocase also functions as an anti-recombinase to displace Rad51 filaments (64,65) from ssDNA (24,66). Once the monomeric translocase removes the RecA filament within a ssDNA gap and reaches the ss/ds DNA junction, the 2B sub-domain duplex DNA interaction would form an inhibited complex and prevent unwanted DNA unwinding beyond the ssDNA gap.

The studies presented here suggest a general mechanism for how dimerization activates the DNA unwinding activity of UvrD-family helicases. However, there is still much that we do not understand about how dimeric SF1A helicases translocate and unwind DNA. For example, even though dimerization relieves 2B-based auto-inhibition, both subunits must still be able to hydrolyze ATP because a hetero-dimer containing a wild type UvrD subunit and an ATPase-deficient subunit is not an active helicase (18,67). We also know that dimerization stimulates the ATPase activity of these enzymes (68-71) and that there is allosteric communication between dimer subunits (67,72). None of these observations are explained by the dimer-dependent re-positioning of the 2B domain away from downstream DNA. The ability to generate 100% covalent dimers that are constitutively active helicases should greatly facilitate further mechanistic studies of these enzymes. The higher processivity of these covalent dimeric helicases may also provide improvements in biotechnological processes, such as isothermal PCR (73,74) and nanopore DNA sequencing (75) that require processive helicases and ssDNA translocases.

## ACKNOWLEDGMENTS

We thank Alex Kozlov and Eric Tomko for critical discussions.

## FUNDING

This work was supported in part by the National Institutes of Health (R35 GM136632 to TML and R35 GM144282 to EAG). Funding for open access charge: National Institute of General Medical Sciences (R35 GM136632).

## Conflict of interest statement

None declared.

## Author Contributions

B.N. and T.M.L. designed research; B.N. an K.N.M. performed experiments and analyzed the data. B.N. and A.C. designed the dimer mutations. B.N., K.N.M, T.M.L. and E.A.G. wrote the paper. T.M.L. and E.A.G. guided the project and provided funding.

## Notes

### Competing Interest Statement

The authors have declared no competing interest.

