## Supplemental Material for "A Conserved Mechanism for Dimerization and Activation of Superfamily 1A UvrD-family Helicases"

**Table S1**: DNA oligomers and their extinction coefficients ($M^{-1}$ $cm^{-1}$ single strands)

| **Sequence abbreviations** | **Number of nucleotides** | **Sequence** | $\varepsilon_{260,\ DNA}$ |
|---|---|---|---|
| 18-TOP-BHQ2 | 18 | GCC CTG CTG CCG ACC AAC -BHQ2 | 156500 |
| 21-TOP-BHQ2 | 21 | GCC CTG CTG CCG ACC AAC GAT -BHQ2 | 188200 |
| 23-TOP-BHQ2 | 23 | GCC CTG CTG CCG ACC AAC GAT GG -BHQ2 | 208600 |
| 25-TOP-BHQ2 | 25 | GCC CTG CTG CCG ACC AAC GAT GGT T -BHQ2 | 225200 |
| 40-TOP-BHQ2 | 40 | GCC CTG CTG CCG ACC AAC GAT GGT TAC ATT CCC GCT GCT G -BHQ2 | 355900 |
| 50-TOP-BHQ2 | 50 | GCC CTG CTG CCG ACC AAC GAT GGT TAC ATT CCC GCT GCT GCT AGT GCA GG -BHQ2 | 452500 |
| CY5-18-BOT-T20 | 38 | CY5- GTT GGT CGG CAG CAG GGC $T_{20}$ | 332200 |
| CY5-21-BOT-T20 | 41 | CY5- ATC GTT GGT CGG CAG CAG GGC $T_{20}$ | 361600 |
| CY5-23-BOT-T20 | 43 | CY5- CCA TCG TTG GTC GGC AGC AGG GC $T_{20}$ | 374600 |
| CY5-25-BOT-T20 | 45 | CY5- AAC CAT CGT TGG TCG GCA GCA GGG C $T_{20}$ | 400400 |
| CY5-40-BOT-T20 | 60 | CY5- CAG CAG CGG GAA TGT AAC CAT CGT TGG TCG GCA GCA GGG C $T_{20}$ | 548800 |
| CY5-50-BOT-T20 | 70 | CY5- CCT GCA CTA GCA GCA GCG GGA ATG TAA CCA TCG TTG GTC GGC AGC AGG GC $T_{20}$ | 638000 |
| HP65 | 65 | GCC TCG CTG CTT TTT GCA GCG AGG C $T_{40}$ | 540900 |

$\varepsilon_{260,\ BHQ2}$ = 8000 $M^{-1}$ $cm^{-1}$; $\varepsilon_{579,\ BHQ2}$ = 38000 $M^{-1}$ $cm^{-1}$

$\varepsilon_{260,\ CY5}$ = 10000 $M^{-1}$ $cm^{-1}$; $\varepsilon_{650,\ CY5}$ = 250000 $M^{-1}$ $cm^{-1}$
